# Thin layer immunoassay; an economical approach to diagnose *Helicobacter pylori* Infection in gastroduodenal ulcer disease patients of Pakistan; a comparative analysis

**DOI:** 10.1101/2020.01.05.895375

**Authors:** Faisal Aziz, Yasmeen Taj, Shahana Urooj Kazmi

## Abstract

*Helicobacter pylori* is a causative agent of gastritis, gastroduodenal ulcers and gastric adenocarcinoma. Gastric patient’s serums were screened for *H. pylori* infection by thin layer immunoassay. A polystyrene plate coated with *H. pylori* sonicate whole cell antigen (10 μg/ml). Two fold-diluted patient’s serum was allowed to react at 37 °C, incubated at 60 °C for 1 min over water bath and recorded water condensation pattern for *H. pylori* antibody. Gastric patient’s blood samples (62% male and 6% female) were tested positive for *H. pylori,* while agewise 15–25 years males (36%) and 65–75 years females (50%) showed highest number of *H. pylori* infection. Thin layer immunoassay showed sensitivity (72–67%), specificity (100%), accuracy (94–69%) and κ value (0.493–0.357) in comparison with wELISA, sELISA and kELISA. We conclude thin layer immunoassay was reliable, low cost, quick, simple and clinically useful method for *H. pylori* diagnosis in patients of Pakistan.

## 1. Introduction

*Helicobacter pylori* is a causative agent of gastritis, ulcer diseases which leads to the development of gastric cancer [1, 2]. It is gram-negative microaerophillic, flagellated and spiral shaped bacilli [3, 4]. More than 50 % world population is colonized by *H. pylori* but few of them suffer from active disease because of various factors, such as; age, gender, crowding, hyperacidity, smoking habit, unhygienic and poor socio-economic status [5, 6, 7]. Developing countries had a prevalence of 80 % and its infection acquired during childhood while the developed countries had a prevalence rate of 20–50 %, which increase in adult life [8, 9]. The developed countries showed age wise prevalence increase at the age between 18 and 29 years (10%); 60–69 of years (27 %); 60–69 years (48 %) while in developing countries prevalence is not linked to the age as 80–90 % prevalence rate all are observed in adult [10]. Persistent infection of *H. pylori* results in the development of local chronic inflammatory and systemic antibody response along with secretion and diffusion of large amount of extracellular products into the mucosa [9]. Major humoral and cellular responses against *H. pylori* infection are unable to eliminate it from host, which results in long lasting infection and constant level of the systemic immune response with high antibody titre in patients. Along with other factors, the improper diagnostic facilities are also responsible for increasing the incidence of *H. pylori* infections [7].

There are number of serological test for the detection of *H. pylori* antibodies like microagglutination assay, enzyme-linked immunosorbent assay (ELISA) and thin layer immunoassay [11]. The test choice should be accordance to the prevalence of the infection, symptoms, practice, time duration, availability and cost [12]. ELISA needs a variety of reagents and sophisticated equipments, but the developing countries have difficulty to arrange such expensive equipment especially at the remote areas of the country [13]. Thin layer immunoassay is quick and simple assay, which does not require sophisticated equipments and carried out by common laboratory equipments and materials at any demographic locations (13, 14, 15). It is sensitive and based on a simple visualization of antigen-antibody reactions by naked eye (16, 17). Because of its simplicity, high capacity and low cost, it can be used for the screening of large number of patient’s serum [15]. The aim of this study was to develop and evaluate thin layer immunoassay for the rapid and economical diagnosis of *H. pylori* infection in patients with gastroduodenal ulcer patients in Pakistan.

## 2. Materials and methods

### 2.1. Place and Duration of Study

It is a descriptive study, was design at the Immunology & Infectious Diseases Research Laboratory, Department of Microbiology, University of Karachi in collaboration with Departments of Medicine and Surgery, Dow University of Health Sciences, Civil Hospital, Karachi, Pakistan, between 2005 to 2009.

### 2.2. Inclusion criteria

All cases coming to the endoscopy unit of Dow University of Health and Sciences, Civil Hospital, Karachi suspected for gastritis and gastroduodenal ulcer diseases on clinical assessment. Cases included children, adults and older persons.

### 2.3. Exclusion criteria

Cases already admitted in the hospital cases on antibiotic treatment.

### 2.4. Patients and gastroendoscopy

All gastric patients were undergone through gastroendoscopy and examination in the Civil Hospital Karachi, Pakistan. Tissue biopsies were obtained from the antral and corpus part of the stomach during gastrointestinal endoscopy along with 214 blood samples from consenting patients. Gastric patient’s blood samples were characterized as gastritis and gastric ulcer. All the samples were transported to Immunology & Infectious Diseases Research Laboratory (IIDRL) for further processes and frozen at −20 °C until tested. All gastric samples collected from the patients and the research protocols were in accordance with the Karachi University’s Ethical Committee.

### 2.6. Preparation of *H. pylori* sonicate whole cell antigen

Tissue biopsies were processed and cultured on Colombia agar (CA) plates containing 7 % lysed horse blood and antibiotics (Amphotericin B, Trimethoprim, Cefsulodin and Vancomycin). The CA plates were incubated for 4–5 days under microaerophillic conditions at 37 °C and identification was carried out by different conventional and molecular methods. Several *H. pylori* strains were inoculated in the BHI broth, incubated at 37 °C for 5 days in microaerophillic environment. *H. pylori* heavy growth was centrifuged at 10,000 rpm for 10 mins, followed by 3 times washing with 20 mM Tris HCl buffer (pH 7.5) and suspended in the same buffer. Cells were sonicated on ice with 6 times for 30 s with a 60 s interval between each shock. The sonicated samples were centrifuged at 5000 rpm for 10 min to remove the cell debris. Then supernatant treated with 0.5 % formalized saline, makes the aliquots, and kept frozen at −20 °C. Protein concentration was measured by Coomassie blue assay (Bio-Rad, USA), bovine serum albumin used as a standard [19].

### 2.7. Thin layer immunoassay

Polystyrene Petri plate was rinsed with 70 % ethanol and dry with a jet of air. BSA solution (20 ml) containing 10 μg/ml of *H. pylori* sonicate whole cell antigen was coated in plate and incubated overnight at 4 °C. Pour off the antigen solution and plate was washed thoroughly gently three times with 150 mM NaCl solution and with distilled water and finally air dried. Draw a grid on the backside of the plate to provide multiple squares. Two-fold serial dilutions of antiserum (1:2 and 1:16384) were prepared in NaC1 (0.15 M) and spot 5 μl antiserum dilution on the respected grid of plate, followed with incubation in humid chamber at 37 °C for 1 h. Plate with serum drops was inverted over a water bath at 60 °C for one minute. Later rinse the plate thoroughly with distilled water, blow dry and again inverted the plate over the water bath at 60 °C for 1 h. Small drops or condensation pattern was regarded as antigen/antibody reaction (Fig. 1). Those dilutions gave water droplets larger than droplets appearing in the background were recorded as positive. Normal control serum was taken as negative control and infected patient serum was taken as positive control. In addition, the optimal concentration and conditions for incubation of the reagents were optimized.

**Figure 1:**
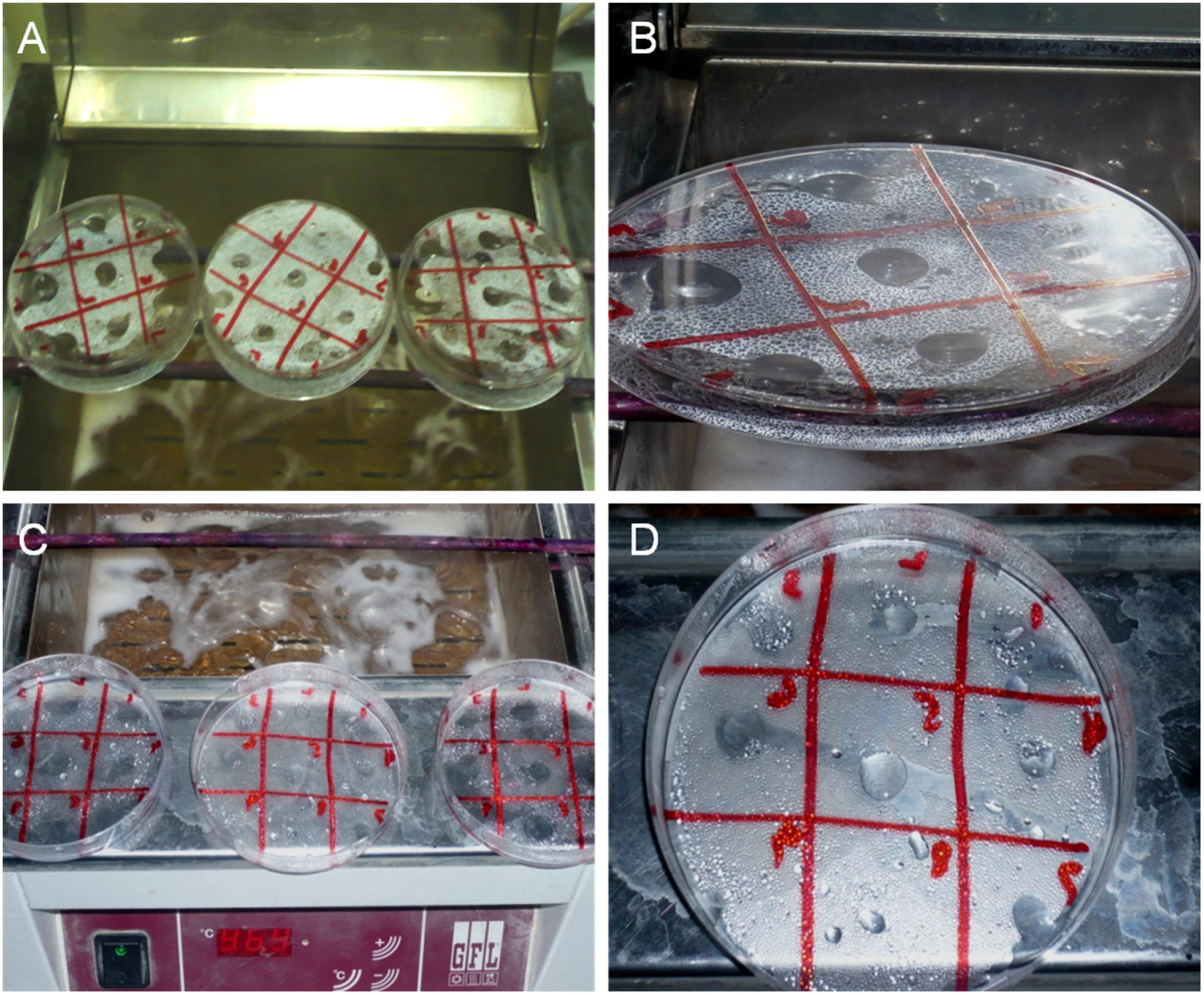
Thin layer immunoassay steps to detect the antibody titre: (a) incubation of polystyrene plate along with different dilution of patient’s serum over water bath. (b) Plate containing droplets after 1 min incubation over bath. (c) Water droplets after rinsing with distilled water, blow dry, and again invert the plate over the water bath at 60 °C. (d) Plates showing water droplets regarded as positive result for *H.pylori*.

### 2.8. Statistical analysis

The serological parameters were optimized and analyzed thin layer immunoassay. The agreement between the thin layer immunoassay and previously reported serological assays as, In-house ELISA based on *H. pylori* surface whole cell antigen (wELISA) [18], In-house ELISA based on *H. pylori* sonicate whole cell antigen (sELISA) [19] and ELISA commercial kit (kELISA) [18] were assessed by the κ statistic.

## 3. Results

This study was designed to develop a thin layer immunoassay for the diagnosis of *H. pylori* infection in gastritis and gastroduodenal patients of Karachi, Pakistan. A total of 214 blood samples were collected from patients reporting at Civil Hospital Karachi with the symptoms of gastritis and gastroduodenal problems. Blood collected from male or female patients (35 % and 65 %, respectively) was processed in IIDRL (Immunology & Infectious Diseases Research Laboratory), Department of Microbiology, Karachi

### 3.1. Analysis of *H. pylori* titer by thin layer immunoassay

A total of 214 human sera samples were screened for anti-*H. pylori* antibodies by thin layer immunoassay. Majority of sera from gastric patients showed low percentage anti-*H. pylori* titre. We found 20 % of patients have 1:128 anti-*H. pylori* titer, while 14 % showed 1:32 and 1:64 titer value. High anti-*H. pylori* titre (1:16384) was found in the 10 % of gastric patients. However, titer of 1:16–1:2 showed 0 % of positive patient, while 2 % samples showed no titre. High titre (1:512 and 1:8192) showed low percentage of positive patients (3 % and 4 %) (Fig. 2).

**Figure 2:**
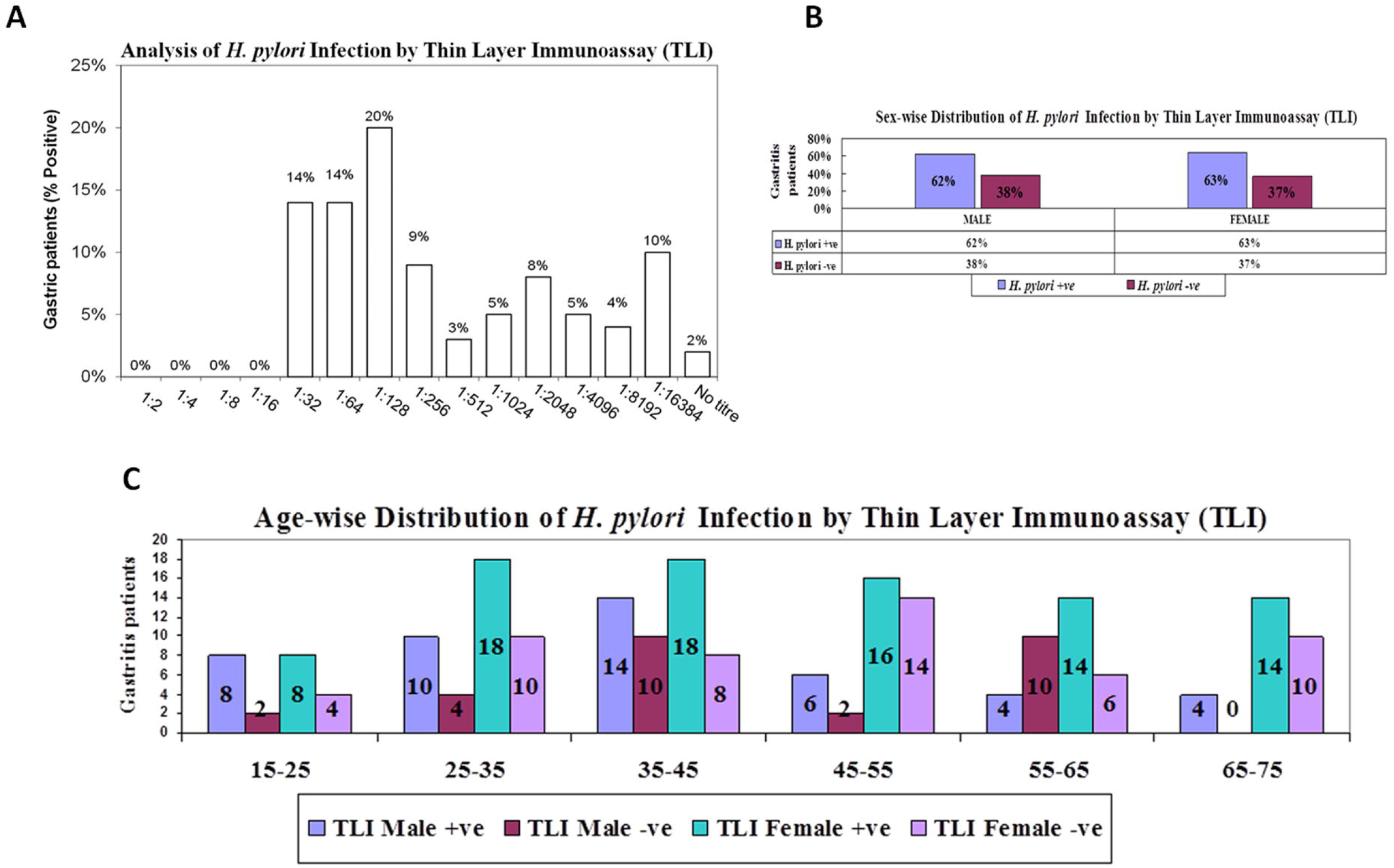
Comparison of anti- *H. pylori* antibody titers between different positive percentages of gastroduodenal ulcer patients by thin layer immunoassay.

### 3.2. Age- and sexwise distribution of gastric patients by thin layer immunoassay

In this study, 214 patients were enrolled between the age of 15 and 75 years (65 % of females and 35 % of males). Overall seropositivity of *H. pylori* infection was found to be 63 % by thin layer immunoassay as compared to 87 % by wELISA [18] and 92 % by sELISA [19]. Female (63 %) and male (62 %) patients were found to be *H. pylori* seropositive with sex-wise distribution (Figure 3). However, age-wise distribution showed slight differences in the risk of *H. pylori* seropositive in gastric patients. We found high risk of infection between low-ages group of 15–25 years (73 %) and 25–35 years (67 %) as compared to middle ages of 45–55 years (58 %) and 55–65 years (53 %) have low risk of infections. Male patients (15–25 years) were shown high risk of infection (36 %), while old-ages patients (55–65 years) were shown low (12 %) prevalence of *H. pylori* infected patients. In contrast, the highest risk of infection in the female population was found between the ages of 65–75 years (50 %). Female patients (15–65 years) were shown a moderate risk of infection (36–43 %) (Fig. 4).

**Figure 3:**
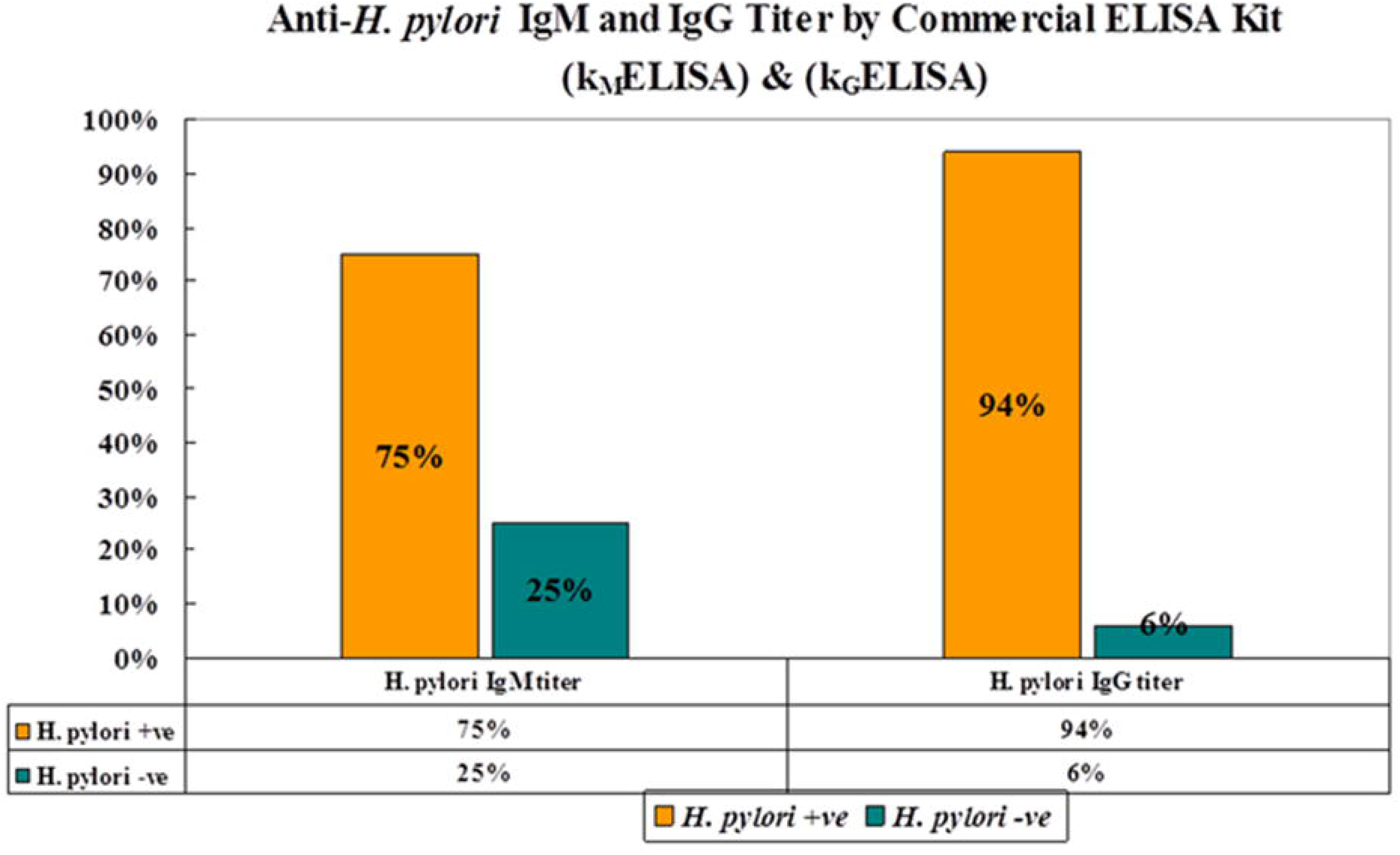
Bar diagram showing high number of male positive result as compare to female by microagglutination method of detection.

**Figure 4:**
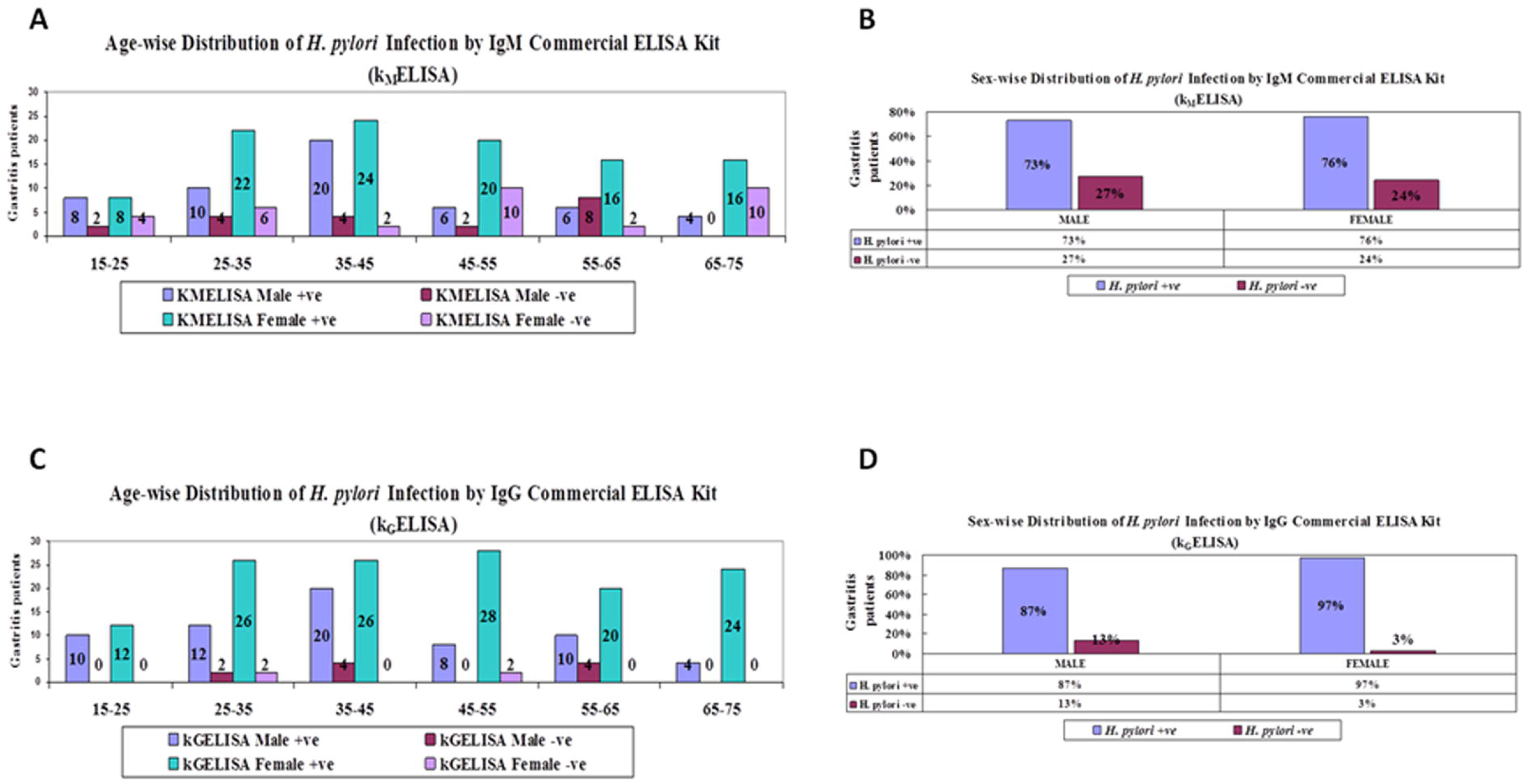
Age wise distribution of *H.pylori* infected gastroduodenal ulcer patients by thin layer immunoassay in reference with sex.

### 3.3. Evaluation of thin layer immunoassay with other serological assays

In order to evaluate the authenticity of thin layer immunoassay; we analyzed the sensitivity, specificity, accuracy and compare with previously reported assay, such as *H. pylori* surface whole cell antigen (wELISA) [18], *H. pylori* sonicate whole cell antigen (sELISA) [19] and commercial ELISA kit (kELISA) [18]. We found moderate agreement of thin layer immunoassay with wELISA (κ value = 0.493) while fair agreement with sELISA (κ value = 0.396) and kELISA (κ value = 0.357). Thin layer immunoassay has sensitivity of 72 % in comparison with wELISA, 68 % with sELISA and 67 % with kELISA. Moreover, we found accuracy of thin layer immunoassay (76 %) in comparison with wELISA, 71 % with sELISA and 69 % with kELISA. Thin layer immunoassay was not observed false positive values while negative predictive values were found in the range of 18–35 %. Based on the accuracy, we can say that the diagnostic value of thin layer immunoassay was to be fair (76 – 71 % = fair test) (Table 1).

## 4. Discussions

*Helicobacter pylori* is a non invasive bacterium which stimulates the immune response by releasing different immunogenic proteins and lipopolysaccharides [20]. ELISA performance is mainly based on the nature of antigen and *H. pylori* strain [21]. *H. pylori* have different type of antigens, such as formaldehyde- or heat-treated whole bacteria, sonic extract, acid glycine extract. Whole cell lysate is the best choice of antigen as compared to a purified antigen which unable to recognize various type of antibodies present in the population. A different population may harbor different natures of *H. pylori* antigen [22]. *H. pylori* local strain antigen is useful with enough influence on the diagnostic properties of serological assay [18].

Serological analysis of *H. pylori* infection is a non invasive and less expensive method [21]. There are many serological assays for *H. pylori* detection which differs on the basis of their sensitivity [7, 12]. These methods have high sensitivities and specificities, yet the expense, time and expertise have lead to a search for an economic, quick and simple serological assay, which is applicable and reliable for developing countries. Thin layer immunoassay is attractive immunoassay in comparison with ELISA because of its rapid result, low cost and simplicity. There is no requirement of sophisticated equipment and standardized reagents, such as enzyme conjugate antibody, substrate and results can be examined by naked eye [13]. Hence, it is appropriate assay to diagnose *H. pylori* infection of gastritis and gastroduodenal ulcer patients from remote areas, where hospital and laboratory’s facility is not well developed. In order to study the seroprevalence of *H. pylori* infection in the population from rural part of Pakistan, we investigated the immune response in gastric patients by developed thin layer immunoassay from the local strain of *H. pylori*. The overall age-wise prevalence rate of *H. pylori* infection was highest among the age group of 15–25 years (73 %) and 25–35 years (67 %) as compared to middle ages of 45–55 years (58 %) and 55–65 years (53 %) have low risk of infections. Male patients (15–25 years) were shown high risk of infection (36 %), while old-ages patients (55–65 years) were shown low (12 %) *H. pylori* infected patients. In contrast, the highest risk of infection in the female population was found between the ages of 65–75 years (50 %). Female patients (15–65 years) were shown a moderate risk of infection (36–43 %) (Fig. 4).

Antibody titer represents the meaningful data to verify the disease status and useful for antibody screening. *H. pylori* antibody is the indication of gastric colonization which may be used as a tool to screen large number of gastric patients. In this study, we found overall moderate prevalence rate (63%) while sex-wise distribution showed little variation as females (63 %) and male (62 %). According to Amjad et al., Pakistanis population has *H. pylori* prevalence of approximately 80.5 % between 1993 and 1997 [7]. In addition, TLI was showed highest titer (1:1,6384) in 22 (10 %) of patients while 42 (20 %) showed 1:128 titer of *H. pylori*. Low numbers of patients 6 (3 %) were showed 1:512 titers.

Thin layer immunoassay based on sonicate whole cell antigen were developed to determine the presence of *H. pylori* infection with a high degree of specificity and sensitivity. We compared thin layer immunoassay with other reported immunoassays that were applied in our laboratory to demonstrate the variability in sensitivity, specificity and other evaluation parameters. Sensitivity can be effect by many factors, such as concentration of reactants, the capacity of the solid phase, the assay speed and incubation temperature. The elevated and long incubation temperature result in the improvement of sensitivity, but the dissociation rate and intra-assay variation can also be high. In comparison of thin layer immunoassay with wELISA [18], sELISA [19] and kELISA [18], we found low sensitivity of 72 %, 68 % and 67 % respectively. However, the accuracy of thin layer immunoassay was in the range of 69–76 % in comparison with wELISA, sELISA and kELISA (Table 1). The moderate association of thin layer immunoassay was found between wELISA (κ value = 0.493), however, fair association with sELISA (κ value = 0.396) and kELISA (κ value = 0.357). Gomez et al., reported high sensitivity and kappa value of thin layer immunoassay compared to ELISA, indicated that thin layer immunoassay can identify more positive samples than ELISA [13]. Ismail et al. reported the comparable sensitivity with low specificity of thin layer immunoassay in comparison with ELISA [14]. According to Nilsson at al. thin layer immunoassay seemed to be less sensitive because of its simplicity and high capacity [15]. False-positive test results may be due to the presence of cross-reacting bacterial antigens [23] that can be reduce by adsorption assay with *H. pylori* closely related species [22]. In our study, we not found false-positive result; hence, no absorption step was considered necessary. The sensitivity, specificity and accuracy indicated thin layer immunoassay as fair test to detect *H. pylori* antibody titer in patients with gastritis and ulcer diseases. Commercial ELISA kits are so expensive and have no surety of high specificity and sensitivity because of the *H. pylori* strain variations. *H. pylori* heterogenicity and cost of commercial ELISA kit limited its application, while in-house serological assays have no such problems [18].

The simplicity, low cost and rapid results of thin layer immunoassay makes its selection as best assay for early diagnosis of *H. pylori* in clinical laboratories, field surveys and rural areas. Furthermore, a large number of samples can be diagnosed because of its rapid screening [24]. Thin layer immunoassay has advantage of giving qualitative as well as quantitative information concerning antibody content. The area recorded is directly related to the logarithmic expression of the antibody concentration [15]. The visualization procedure of thin layer immunoassay is one of its advantage that an antigen/antibody reaction may be detected by naked eye, based on the principal of vapor condensation indicated an area of antigen/antibody reaction, contain high protein and more hydrophilic than the remaining surrounding reaction (25, 24). The result of thin layer immunoassay can be saved by preserve polystyrene plate and this storage did not cause any false positive or false-negative results, even though there was an apparent growth of microorganism in the samples [25]. However, thin layer immunoassay has the disadvantage of requiring a higher concentration of antigens and can’t be applied for high antibody titre detection [14].

## 5. Conclusions

An ideal serodiagnostic test, especially for developing countries, should be low cost, rapid and easy to perform, in addition to being sensitive and specific. Thin layer immunoassay fulfills these requirements and suitable to be used in field studies to get medical attention on right time, as required in outbreaks of infection. We conclude, thin layer immunoassay is reliable, low cost, simple and clinical useful assay to diagnosis *H. pylori* infection in gastric patients of Pakistan.

## Table Caption

Table 1: Table showing comparative analysis of thin layer immunoassay with other comparative serological assays [18, 19].

## Acknowledgments

Financial disclosures funds to conduct study were provided by Higher Education Commission (HEC), Pakistan. No other financial declarations by any of the authors.

## Abbreviations

ELISA: Enzyme-linked immunosorbent assay
*H. pylori*: *Helicobacter pylori*
kELISA: ELISA commercial kit
OD: Optical density
wELISA: In-house ELISA based on *H. pylori* surface whole cell antigen
sELISA: In-house ELISA based on *H. pylori* sonicate whole cell antigen

