## Supplementary material for "Thin layer immunoassay; an economical approach to diagnose *Helicobacter pylori* Infection in gastroduodenal ulcer disease patients of Pakistan; a comparative analysis": table1

**Table 1: Anti- *H. pylori* IgM antibody titer values in gastroduodenal ulcer patients by commercial kit ELISA (k_M_ELISA)**

| Negative controls | 0.015 | Positive controls | 1.405 |
| --- | --- | --- | --- |
|  | 0.014 |  | 1.157 |
|  | 0.0106 |  | 1.074 |
| Mean ± SD | 0.013 ± 0.002 | Mean ± SD | 1.212 ± 0.172 |
| Cutoff value = Mean of negative control + 0.250 | | | 0.263 |
| Prevalence of *H. pylori* by IgM ELISA kit | | | 75 % (0.921 ± 0.633) |
