## Supplementary material for "Thin layer immunoassay; an economical approach to diagnose *Helicobacter pylori* Infection in gastroduodenal ulcer disease patients of Pakistan; a comparative analysis": table2

**Table 2:** Anti- *H. pylori* IgG antibody titer values in gastroduodenal ulcer patients by commercial kit ELISA (k_G_ELISA)

| Negative controls | 0.0037 | Positive controls | 1.006 |
| --- | --- | --- | --- |
|  | 0.0032 |  | 1.179 |
|  | 0.0038 |  | 1.083 |
| Mean ± SD | 0.003 ± 0.0003 | Mean ± SD | 1.089 ± 0.0866 |
| Cutoff value = Mean of negative control + 0.250 | | | 0.253 |
| Prevalence of *H. pylori* by IgG ELISA kit | | | 94 % (2.144 ± 0.997) |
