## Supplementary material for "Thin layer immunoassay; an economical approach to diagnose *Helicobacter pylori* Infection in gastroduodenal ulcer disease patients of Pakistan; a comparative analysis": table3

Table 3: Table showing statistical analysis of thin layer immunoassay with previously reported serological assays.

| Standard Assays | Sensitivity  % | Specificity  % | Accuracy  % | PPV  % | NPV  % | FPV | FNV | OA | EA | AABC | PABC | KAPPA |
| --- | --- | --- | --- | --- | --- | --- | --- | --- | --- | --- | --- | --- |
|  | *H. pylori* thin layer-immunoassay | | | | | | | | | | | |
| wELISA | 72 | 100 | 76 | 100 | 35 | 0.0 | 0.28 | 0.757 | 0.52 | 0.237 | 0.480 | 0.493 |
| sELISA | 68 | 100 | 71 | 100 | 22.5 | 0.0 | 0.316 | 0.710 | 0.52 | 0.190 | 0.480 | 0.396 |
| kELISA | 67 | 100 | 69 | 100 | 18 | 0.0 | 0.33 | 0.691 | 0.52 | 0.17 | 0.480 | 0.357 |
